# A brainstem circuit for gravity-guided vertical navigation

**DOI:** 10.1101/2024.03.12.584680

**Authors:** Yunlu Zhu, Hannah Gelnaw, Franziska Auer, Kyla R. Hamling, David E. Ehrlich, David Schoppik

## Abstract

The sensation of gravity anchors our perception of the environment and is crucial for navigation. However, the neural circuits that transform gravity into commands for navigation are undefined. We first determined that larval zebrafish (*Danio rerio*) navigate vertically by maintaining a consistent heading across a series of upward climb or downward dive bouts. Gravity-blind mutant fish swim with more variable heading and excessive veering, leading to inefficient vertical navigation. After targeted photoablation of ascending vestibular neurons and spinal projecting midbrain neurons, but not vestibulospinal neurons, vertical navigation was impaired. These data define a sensorimotor circuit that uses evolutionarily-conserved brainstem architecture to transform gravitational signals into persistent heading for vertical navigation. The work lays a foundation to understand how vestibular inputs allow animals to move efficiently through their environment.

## INTRODUCTION

Animals adopt navigational strategies tailored to their sensory ecology^1,2^. Perception of the environment, particularly for species that swim or fly^3–6^, is anchored by the sense of gravity^7,8^. Vertebrates use otolithic organs in the vestibular system of the inner ear to transduce linear acceleration due to gravity^9^. Vestibular information has long been thought to impact spatial navigation^10^, shaping behaviors such as stabilization of vision and posture, perception of self-motion and head direction, motor coordination, and path integration^8,11–13^, and can modulate the activity of neurons responsible for navigation, such as head direction cells of the mammalian limbic system^14–16^. However, complete circuits that transform sensed gravity into motor signals for navigation remain undefined.

The larval zebrafish (*Danio rerio*), a small translucent vertebrate, is an ideal model to discover neural substrates for gravity-guided navigation^17–21^. Zebrafish larvae translate in short discrete bouts punctuated by periods of inactivity. The separable active and passive phases of locomotion facilitates dissection of the neural-derived commands for movement from their biomechanical consequences. Zebrafish maintain a dorsal-up orientation relative to gravity using postural reflexes^22–26^ and learned control of movement timing^27^. These vestibular behaviors rely entirely on otolithic organs^28^, in particular the gravity-sensing utricle.^29–31^. Anatomically, utricle-recipient vestibular nuclei relay information to the spinal cord directly^32–34^ and indirectly^24,25,30,35,36^ through a highly-conserved midbrain population called the interstitial nucleus of Cajal / nucleus of the medial longitudinal fasciculus (INC/nMLF)^37,38^. Finally, while zebrafish navigate in the horizontal plane^39–41^ it is unclear if they similarly maintain vertical heading to navigate in depth.

Here, we combined high-throughput behavioral analysis of vertical locomotion and loss-of-function assays to explore neural circuits for gravity-guided navigation. We first established that larvae swim in a series of bouts with consistent heading to navigate in the dark. Stable control of heading allowed larvae to efficiently change depth. Gravity sensation is essential for this navigation behavior, as mutant fish without utricular otoliths navigate depth poorly, swimming with more variable heading and excessive veering. Lesions of ascending utricle-recipient neurons in the tangential vestibular nucleus recapitulated this phenotype, while lesions of descending vestibulospinal neurons did not. The INC/nMLF receives ascending inputs; lesions there disrupted heading and navigation efficacy. Taken together, our data reveals a conserved hindbrain-midbrain-spinal cord circuit that transformed sensed gravity to commands to maintain heading for effective vertical navigation. More broadly, we reveal ancient architecture that leverages sensed gravity to move efficiently through the world.

## RESULTS

### Larval zebrafish navigate depth by maintaining a consistent heading over a series of swim bouts

We first examined whether larval zebrafish maintain a consistent heading as they navigate in depth^42^. To measure behavior, we used a high-throughput real time Scalable Apparatus to Measure Posture and Locomotion (SAMPL)^43^. SAMPL records body position and posture in the pitch axis (nose-up/nose-down) as larval zebrafish swim freely in depth (Figure 1A). We examined freely swimming larvae from 7 to 9 days post-fertilization (dpf) in complete darkness. We measured the trajectory of swim directions relative to horizontal and observed both upward and downward swim bouts (Figures 1B and 1C), indicating that larvae climb and dive in the water column. To quantify the spread of swim directions, we defined variability as the median absolute deviation of swim bouts (Figure 1D).

**Figure 1:**
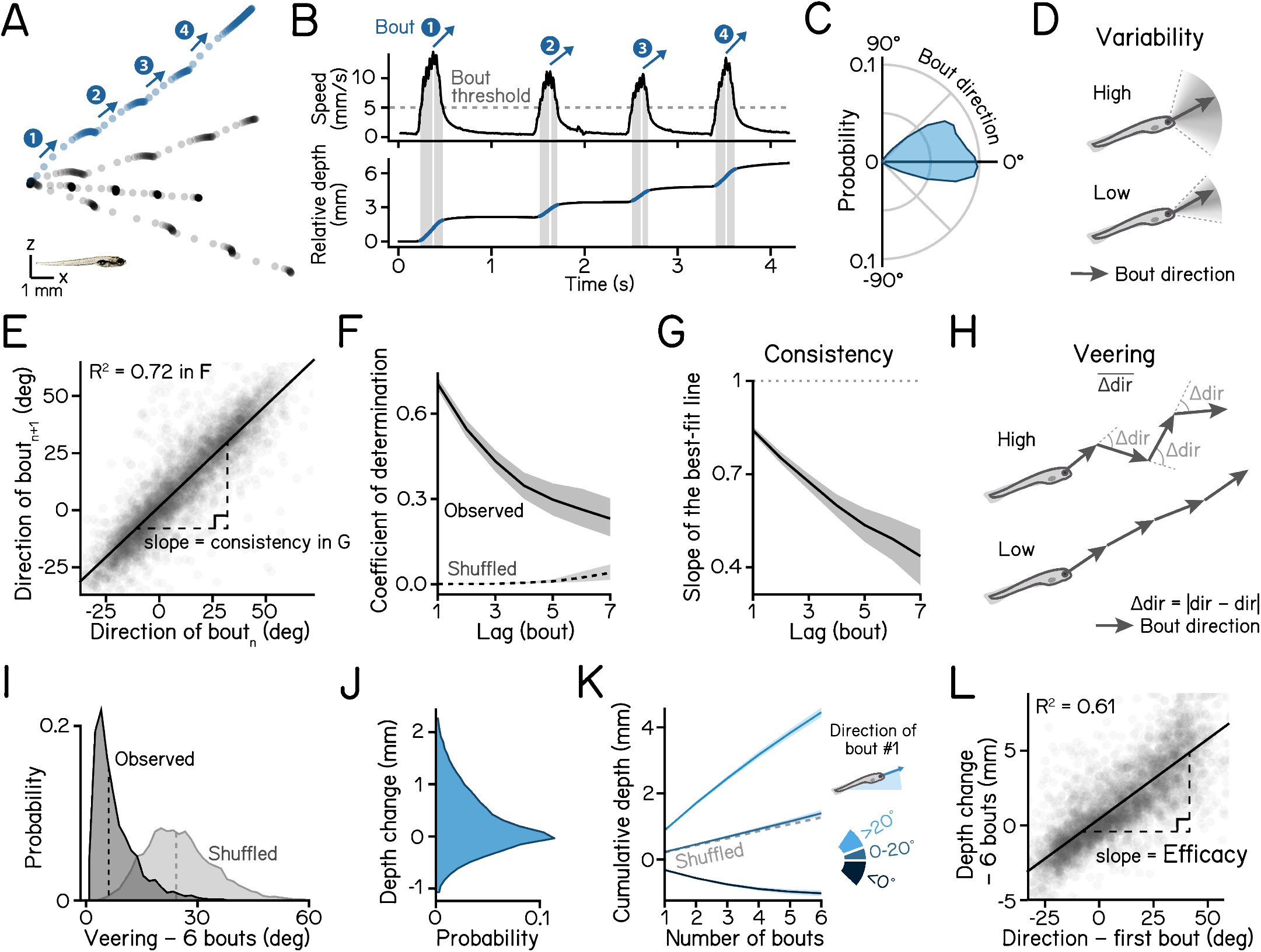
Larvae navigate depth in series of consecutive bouts with consistent heading. **(A)** Sample swim trajectories of 7 dpf larvae in the x/z axes. All trajectories begin from the left. Dots represent fish locations 48 ms apart (down-sampled from 166 Hz data for visualization). Arrows mark swim directions of individual bouts in the blue trajectory. Scale bar: 1 mm. **(B)** Time series data of the blue trajectory in A. Horizontal dashed line in the upper panel indicates the 5 mm/s threshold for bout detection. Vertical lines label the time of peak speed for each bout. Lower panel plots directions of movement (black) and body posture in the pitch axis (orange). **(C)** Polarized histograms (frequency polygons) of bout directions of three WT zebrafish strains. n = 121,979 bouts from 537 fish. **(D)** Schematic illustrations of bout direction variability. A wide distribution of bout directions indicates high variability. **(E)** Directions of the following bout plotted as a function of the current plot. Correlation coefficient is plotted in (F). Slope of the best fitted line is plotted in (G). n = 61,990 bout pairs from 537 fish. **(F)** Serial correlation (autocorrelation) of swim directions across observed consecutive bouts and shuffled bouts. 95% correlation confidence intervals are shown as shaded error bands. **(G)** Slope of the best fit line of swim directions of bout(n+lag) vs. bout(n) is defined as the swim direction consistency. 95% confidence intervals of the estimated slope are shown as shaded bands. **(H)** Veering is quantified as the absolute change of swim directions between adjacent bouts, averaged through a bout series. A course of trajectory with greater direction changes results in higher veering. **(I)** Veering across 6 consecutive bouts (observed) and shuffled bouts are plotted as histograms. Median values are shown as dashed lines. n = 4,048 sets of 6 bouts. *P*_*median-test*_ < 1e-16. **(J)** Distribution of depth changes, defined as the displacement on the z axis, of a single swim bout. n = 121,979 bouts from 537 fish. **(K)** Cumulative depth change through 6 consecutive bouts, separated by the swim direction of the first bout. **(L)** Depth change efficacy through 6 bouts, defined as the slope of best fit line of total depth change during the bout series vs. swim direction of the first bout. n = 4,048 sets of 6 bouts. See also Table 1 for parameter definitions and statistics.

The depth change resulting from a single swim bout was small (0.34 mm, median of absolute depth displacement), so we hypothesized that larval zebrafish integrate a series of swim bouts to adjust their depth efficiently. We quantified and parameterized the statistics of short series of sequential bouts. Directions of consecutive bouts were highly correlated (Figure 1E), determined by the coefficient of determination of direction (Figure 1F), and highly consistent, defined as the slope of the best-fit line between directions of consecutive bout (Figure 1G, and Table 1 for parameter definitions and statistics). As the series continued, bout direction became increasingly less correlated with the first bout (Figures 1F and 1G). To quantify the amount of direction change during consecutive bouts, we defined veering as the mean of absolute direction differences between adjacent bouts (Figure 1H). Compared to shuffled bouts, fish veered significantly less during observed consecutive bouts (Figure 1I, 6.08 deg vs. 23.90 deg, observed vs. shuffled, *P*_*median-test*_ < .001), indicating that larval zebrafish maintain stable swim directions through a series. Consequentially, a bout series results in cumulative changes in depth (Figure 1K). The cumulative depth change across a series of bouts is highly correlated with the direction of the first swim bout in the series (Figure 1J). We therefore defined the efficacy of depth change as the slope of the best fitted line between cumulative depth change and the direction of the first bout in the sequence (Figure 1L). Given the swim direction of a bout, a higher efficacy represents a greater depth change achieved through following consecutive bouts in the sequence.

**Table 1:** Parameters definitions and statistics. Refer to Figure 1. Definitions of navigation parameters and values of wild-type 7-day larvae. Values that can be calculated without regression analysis are reported as median across all swim bouts (n = 121957 bouts) or across 6 consecutive bouts (n = 4047). Regression coefficients such as efficacy and consistency are reported as mean fitted slope across all experimental repeats (N = 27). Bout kinematics are shown as mean across experimental repeats (N = 27).

| Parameter | Definition | Format | Value | Unit |
| --- | --- | --- | --- | --- |
| Swim bout | An epoch during which fish swims faster than 5 mm/s | – | – | – |
| Swim direction | Swim direction at the time of peak speed | Median [IQR] | 9.01 [33.76] | deg |
| Direction variability | Median absolute deviation (MAD) of all swim bout directions | MAD | 16.77 | deg |
| Veering | Absolute differences of directions between adjacent bouts averaged across a series of 6 consecutive bouts | Median [IQR] | 6.08 [6.79] | deg |
| Depth change | Displacement on the vertical axis during a swim bout | Median [IQR] | 0.14 [0.75] | mm |
| Absolute depth change | Distance on the vertical axis during a swim bout | Median [IQR] | 0.34 [0.56] | mm |
| Cumulative depth change | Cumulative displacement in depth through 6 consecutive bouts | Median [IQR] | 0.46 [1.64] | mm |
| Depth change efficacy | Slope of the best fit line of cumulative depth change vs direction of the first bout in the sequence | Mean [SD] | 9.91e-2 [1.76e-2] | mm/deg |
| Direction consistency (lag = 1) | Slope of the best fit line of direction of bout(n+lag) vs that of bout(n) | Mean [SD] | 0.84 [3.07e-2] | – |
| Direction consistency (lag = 2) | – | Mean [SD] | 0.75 [4.78e-2] | – |
| Direction consistency (lag = 3) | – | Mean [SD] | 0.68 [7.04e-2] | – |
| Direction consistency (lag = 4) | – | Mean [SD] | 0.60 [9.51e-2] | – |
| Direction consistency (lag = 5) | – | Mean [SD] | 0.54 [0.14] | – |
| Direction consistency (lag = 6) | – | Mean [SD] | 0.49 [0.18] | – |
| Direction consistency (lag = 7) | – | Mean [SD] | 0.44 [0.22] | – |
| Steering gain | Slope of the best fit line of direction vs posture at time of the peak speed | Mean [SD] | 0.68 [3.20e-2] | – |
| Lifting gain | Slope of the best fit line of estimated lift vs bout depth change | Mean [SD] | 0.36 [3.80e-2] | – |
| Righting gain | Absolute value of the slope of the best fit line of rotation during deceleration vs initial posture | Mean [SD] | 0.18 [1.79e-2] | – |

We conclude that larvae change depth efficiently by performing a series of swim bouts with consistent heading. The parameters of consistency, veering, and efficacy define their ability to navigate in depth.

### Loss of gravity sensation disrupts vertical navigation

To understand whether the sensation of gravity contributes to navigation in depth, we examined behavior of 7–9 dpf gravity-blind larvae as they swam in complete darkness. The *otogelin* mutant fails to develop a utricular otolith (Figure 2A, arrowhead) until ∼14 dpf^44^, leaving larvae unable to sense gravity.^29,30,45^. Compared to heterozygous siblings, *otog*-/- larvae showed more variable swim directions (Figure 2B. 20.44 deg vs. 21.36 deg, heterozygous controls vs mutants, *P*_*bootstrap*_ = .004, Table 2). In addition, series of bouts by mutants exhibited lower direction consistency (Figure 2C), and veered more (Figure 2D, 5.33 deg vs. 6.21 deg, *P*_*median-test*_ = .006). Consequentially, gravity-blind fish were dramatically less efficient at navigating in depth (Figure 2E, 0.14 deg vs. 9.05e-2 deg, *P*_*bootstrap*_ < .001), in congruence with their high veering. We conclude that gravity sensation is crucial to stabilize heading for efficient vertical navigation.

**Table 2:** Effects of vestibular impairments on locomotion parameters. Refer to Figure 2-4 & S2-S3. Methods of statistical analysis are reported with P values. All P values are two-tailed.

| Parameter | Format | Control value | Condition value | P value | Notes |
| --- | --- | --- | --- | --- | --- |
| **otog mutation** 99/136 fish for hets/mutants |  |  |  |  |  |
| Variability (deg) | Mean [SD] | 20.44 [0.18] | 21.36 [0.26] | $P_{bootstrap} = 4.20e-3$ | Bootstrapped mean and SD |
| Veering (deg) | Median [95CI] | 5.33 [4.90-5.80] | 6.21 [5.90-6.57] | $P_{median-test} = 6.41e-3$ | - |
| Efficacy (mm/deg) | Mean [SD] | 0.14 [3.80e-3] | 9.05e-2 [1.93e-3] | $P_{bootstrap} = 1.12e-27$ | Bootstrapped mean and SD |
| Steering Gain | Mean [SEM] | 0.69 [1.01e-2] | 0.85 [1.52e-2] | $P_{t-test} = 2.33e-5$ | Unpaired t-test |
| Lifting Gain | Mean [SEM] | 0.32 [1.45e-2] | 0.15 [2.95e-2] | $P_{t-test} = 7.09e-4$ | Unpaired t-test |
| Righting Gain | Mean [SEM] | 0.17 [1.08e-2] | 0.09 [1.03e-2] | $P_{t-test} = 6.76e-4$ | Unpaired t-test |
| Consistency (lag = 1) | Mean [SD] | 0.87 [6.90e-3] | 0.87 [5.39e-3] | $P_{bootstrap} = .930$ | Bootstrapped mean and SD |
| Consistency (lag = 2) | Mean [SD] | 0.79 [1.27e-2] | 0.76 [8.02e-3] | $P_{bootstrap} = 4.76e-2$ | - |
| Consistency (lag = 3) | Mean [SD] | 0.73 [1.79e-2] | 0.67 [1.11e-2] | $P_{bootstrap} = 8.56e-3$ | - |
| Consistency (lag = 4) | Mean [SD] | 0.66 [2.52e-2] | 0.57 [1.86e-2] | $P_{bootstrap} = 7.74e-3$ | - |
| Consistency (lag = 5) | Mean [SD] | 0.60 [2.99e-2] | 0.50 [1.90e-2] | $P_{bootstrap} = 2.01e-2$ | - |
| **Tangential lesions** 40/25 fish for controls/lesions |  |  |  |  |  |
| Variability (deg) | Mean [SD] | 20.87 [0.15] | 22.07 [0.23] | $P_{bootstrap} = 8.62e-6$ | Bootstrapped mean and SD |
| Veering (deg) | Median [95CI] | 6.79 [6.48-7.06] | 7.23 [7.02-7.61] | $P_{median-test} = 1.44e-2$ | - |
| Efficacy (mm/deg) | Mean [SD] | 0.10 [2.09e-3] | 8.84e-2 [2.22e-3] | $P_{bootstrap} = 1.87e-4$ | Bootstrapped mean and SD |
| Steering Gain | Mean [SEM] | 0.70 [1.98e-2] | 0.80 [2.23e-2] | $P_{t-test} = 7.41e-3$ | Unpaired t-test |
| Lifting Gain | Mean [SEM] | 0.31 [2.88e-2] | 0.20 [4.02e-1] | $P_{t-test} = 3.57e-2$ | Unpaired t-test |
| Righting Gain | Mean [SEM] | 0.14 [6.59e-3] | 0.13 [9.73e-3] | $P_{t-test} = .180$ | Unpaired t-test |
| Consistency (lag = 1) | Mean [SD] | 0.88 [5.75e-3] | 0.86 [6.23e-3] | $P_{bootstrap} = 9.08e-2$ | Bootstrapped mean and SD |
| Consistency (lag = 2) | Mean [SD] | 0.82 [9.10e-3] | 0.78 [8.16e-3] | $P_{bootstrap} = 1.31e-2$ | - |
| Consistency (lag = 3) | Mean [SD] | 0.78 [1.26e-2] | 0.72 [1.62] | $P_{bootstrap} = 4.53e-3$ | - |
| Consistency (lag = 4) | Mean [SD] | 0.76 [1.00e-2] | 0.69 [1.87] | $P_{bootstrap} = 3.61e-2$ | - |
| Consistency (lag = 5) | Mean [SD] | 0.76 [2.05e-2] | 0.66 [2.37] | $P_{bootstrap} = 7.82e-3$ | - |
| **Vestibulospinal lesions** 79/97 fish for controls/lesions |  |  |  |  |  |
| Variability (deg) | Mean [SD] | 19.87 [0.16] | 24.29 [0.20] | $P_{bootstrap} = 3.87e-70$ | Bootstrapped mean and SD |
| Veering (deg) | Median [95CI] | 10.68 [10.22-11.28] | 9.84 [9.39-10.31] | $P_{median-test} = 1.88e-2$ | - |
| Efficacy (mm/deg) | Mean [SD] | 5.64e-2 [3.18e-3] | 0.10 [2.41e-3] | $P_{bootstrap} = 1.17e-27$ | Bootstrapped mean and SD |
| Steering Gain | Mean [SEM] | 0.60 [2.27e-2] | 0.75 [3.40e-2] | $P_{t-test} = 2.32e-3$ | Unpaired t-test |
| Lifting Gain | Mean [SEM] | 0.41 [2.74e-2] | 0.27 [3.16e-2] | $P_{t-test} = 5.90e-3$ | Unpaired t-test |
| Righting Gain | Mean [SEM] | 0.17 [5.38e-3] | 0.11 [1.22e-2] | $P_{t-test} = 1.00e-3$ | Unpaired t-test |
| Consistency (lag = 1) | Mean [SD] | 0.70 [4.70e-3] | 0.82 [4.73e-3] | $P_{bootstrap} = 2.83e-26$ | Bootstrapped mean and SD |
| Consistency (lag = 2) | Mean [SD] | 0.60 [1.23e-2] | 0.74 [8.95e-3] | $P_{bootstrap} = 2.02e-20$ | - |
| Consistency (lag = 3) | Mean [SD] | 0.54 [1.86e-2] | 0.70 [1.26e-2] | $P_{bootstrap} = 2.89e-14$ | - |
| Consistency (lag = 4) | Mean [SD] | 0.49 [2.39e-2] | 0.66 [2.49e-2] | $P_{bootstrap} = 1.02e-6$ | - |
| Consistency (lag = 5) | Mean [SD] | 0.45 [2.40e-2] | 0.64 [2.85e-2] | $P_{bootstrap} = 1.25e-7$ | - |
| **INC/nMLF lesions** 37/36 fish for controls/lesions |  |  |  |  |  |
| Variability (deg) | Mean [SD] | 14.29 [0.12] | 14.60 [0.12] | $P_{bootstrap} = 7.53e-2$ | Bootstrapped mean and SD |
| Veering (deg) | Median [95CI] | 5.98 [5.74-6.18] | 6.51 [6.27-6.87] | $P_{median-test} = 2.13e-3$ | - |
| Efficacy (mm/deg) | Mean [SD] | 0.14 [3.46e-3] | 0.11 [2.66e-3] | $P_{bootstrap} = 3.00e-13$ | Bootstrapped mean and SD |
| Steering Gain | Mean [SEM] | 0.68 [1.37e-2] | 0.70 [1.89e-2] | $P_{t-test} = .357$ | Unpaired t-test |
| Lifting Gain | Mean [SEM] | 0.34 [2.00e-2] | 0.33 [1.43e-2] | $P_{t-test} = .761$ | Unpaired t-test |
| Righting Gain | Mean [SEM] | 0.21 [4.28e-3] | 0.18 [1.01e-2] | $P_{t-test} = 2.95e-2$ | Unpaired t-test |
| Consistency (lag = 1) | Mean [SD] | 0.84 [6.11e-3] | 0.82 [5.30e-3] | $P_{bootstrap} = 3.31e-3$ | Bootstrapped mean and SD |
| Consistency (lag = 2) | Mean [SD] | 0.75 [6.55e-3] | 0.72 [7.30e-3] | $P_{bootstrap} = 8.52e-2$ | - |
| Consistency (lag = 3) | Mean [SD] | 0.69 [9.48e-3] | 0.63 [1.31e-2] | $P_{bootstrap} = 5.40e-3$ | - |
| Consistency (lag = 4) | Mean [SD] | 0.60 [2.69e-2] | 0.55 [2.53e-2] | $P_{bootstrap} = 8.01e-2$ | - |
| Consistency (lag = 5) | Mean [SD] | 0.58 [3.03e-2] | 0.52 [2.41e-2] | $P_{bootstrap} = .122$ | - |

**Figure 2:**
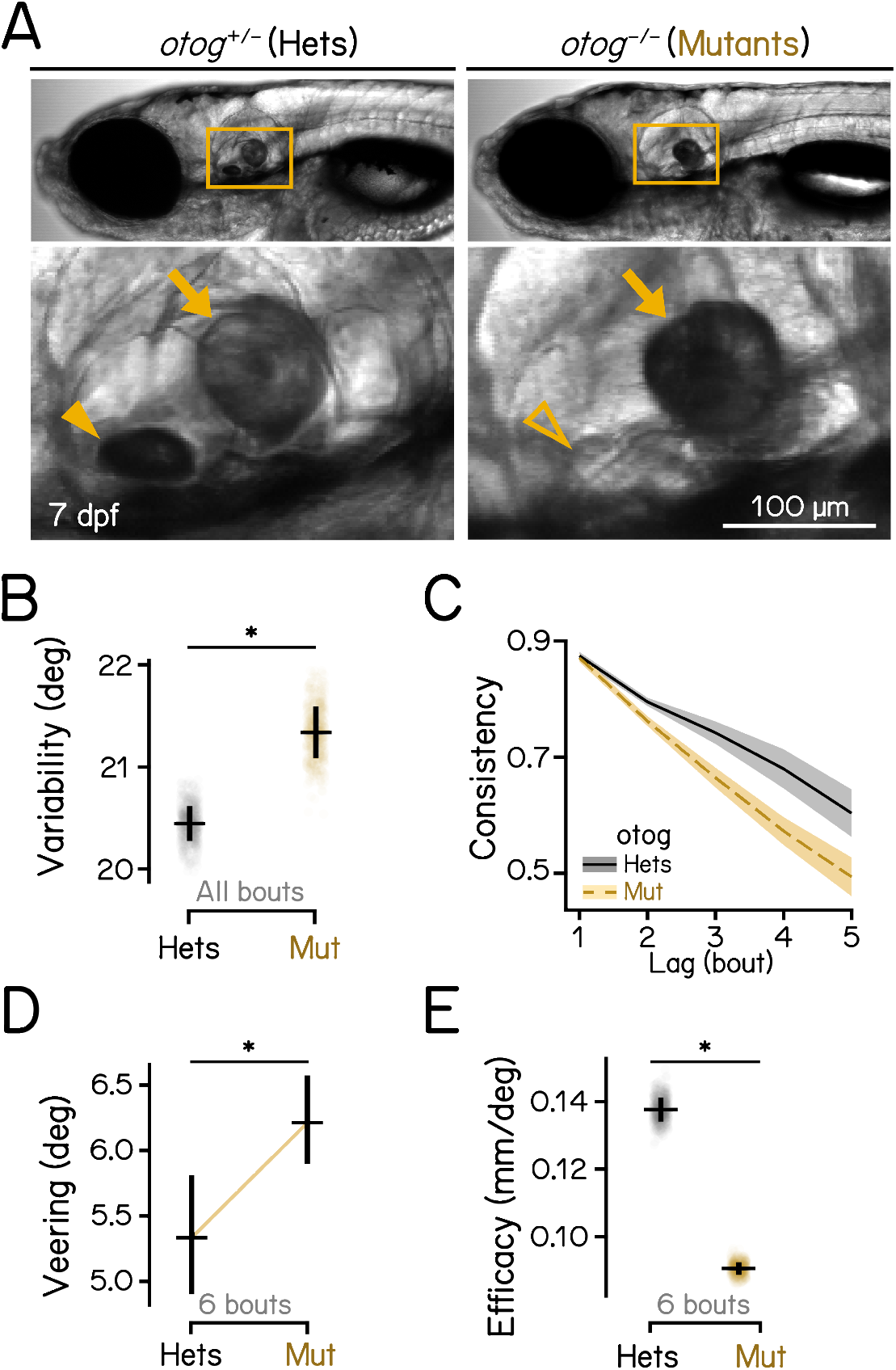
Gravity-blind mutant fish have impaired vertical navigation. **(A)** Homozygous *otog* mutants have intact saccular otoliths (arrows) while lack the utricular otoliths (arrowheads) at 7 dpf. Scale bar: 100 µm. **(B)** Swim direction variability, quantified as the median absolute deviation (MAD) of swim directions of all bouts, compared between *otog* mutants and heterozygous controls. Bootstrapped MAD are plotted as data points with error bars showing standard deviations. n = 14590/10645 bouts from 99/136 fish for controls/mutants. *P*_*bootstrap*_ = 4.20e-3. **(C)** Swim direction consistency, as defined in Figure 1G, plotted as a function of the number of bouts in the sequence. Shaded bands indicate standard deviations of the slope estimated using bootstrapping. **(D)** Veering through 6 consecutive bouts, as defined in Figure 1H, compared between *otog* mutants and heterozygous controls. Median with 95% confidence intervals are plotted. n = 443/1339 6-bout series for controls/mutants. *P*_*median-test*_ = 6.41e-3. **(E)** Depth change efficacy, as defined in Figure 1L. Bootstrapped slopes are plotted as data points with error bars showing standard deviations. *P*_*bootstrap*_ = 1.12e-27. See also Table 2 for statistics.

### Ablation of gravity-sensitive vestibular neurons disrupts vertical navigation and swim kinematics

Previous studies demonstrate that gravity-sensitive ascending neurons of the the tangential vestibular nucleus^46^ and descending vestibulospinal neurons of the lateral vestibular nucleus^47^ encode body tilt and regulate postural behaviors^30,33–36,48^. We adopted a loss-of-function approach, using a pulsed infrared laser to ablate genetically-defined populations of ascending neurons in the tangential nucleus (Figure S1A). In addition, we reanalyzed a dataset^34^ comprised of larvae with lesioned descending vestibulospinal neurons (Figure S1B).

Loss of ascending neurons in the tangential nucleus (Figures 3A and 3B) recapitulated disruption to vertical navigation seen in gravity-blind fish. After lesions, fish had more variable swim directions (Figure 3C; 20.87 deg vs. 22.07 deg, control vs. lesions, *P*_*bootstrap*_ < .001. See also Table 2). Heading consistency was reduced (Figure 3D), and veering increased (Figure 3E, 6.79 deg vs. 7.23 deg, *P*_*median-test*_ = .014), disrupting depth change efficiency (Figure 3F, 0.10 deg vs. 8.84e-2 deg, *P*_*bootstrap*_ < .001).

**Figure 3:**
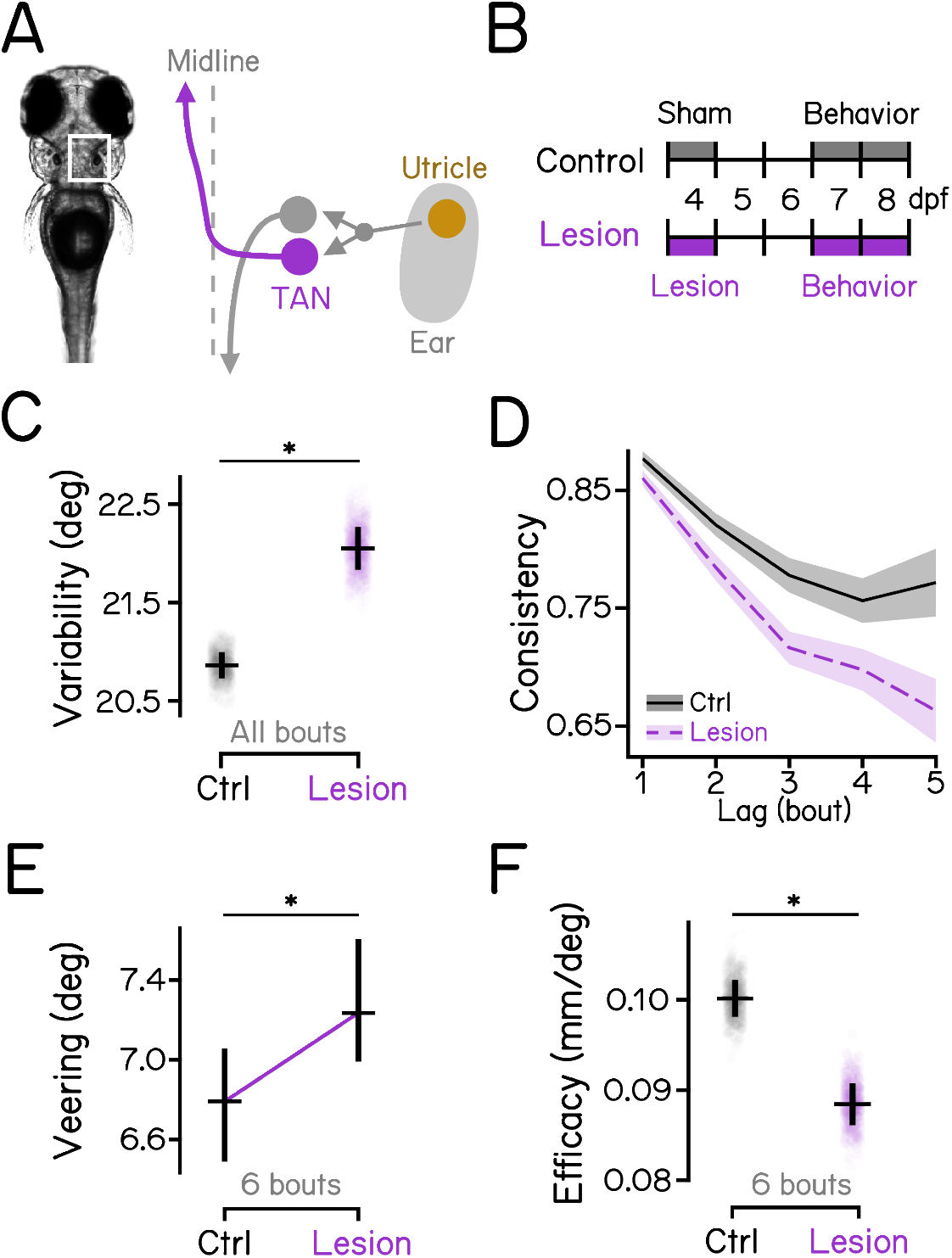
Ascending neurons in the tangential nucleus are indispensable for vertical navigation. **(A)** Schematic view of the inner-ear utricular otolith and the vestibular pathways in the hindbrain of zebrafish. Utricle: utricular otoliths (yellow); TAN: the tangential vestibular nucleus (magenta). **(B)** Diagrams of experimental procedures for lesions of the tangential nucleus and behavioral assays. See Figure S1A for examples of lesions. **(C)** Swim direction variability compared between tangential-lesioned larvae and controls. Bootstrapped MAD are plotted as data points with error bars indicating standard deviations. n = 17797/11417 bouts from 40/25 fish for controls/lesions. *P*_*bootstrap*_ = 8.62e-6. **(D)** Swim direction consistency plotted as a function of the number of bouts in the sequence. Shaded bands indicate standard deviations of the slope estimated using bootstrapping. **(E)** Veering through 6 consecutive bouts plotted in median with 95% confidence intervals. n = 1384/892 6-bout series for controls/lesions. *P*_*median-test*_ = 1.44e-2. **(F)** Depth change efficacy plotted with bootstrapped slopes plotted as data points and error bars showing standard deviations. *P*_*bootstrap*_ = 1.87e-4. See also Table 2 for statistics.

Similar to loss of ascending neurons, lesions of vestibulospinal neurons (Figures S2A and S2B) also increased swim direction variability (Figure S2C; 19.87 deg vs. 24.29 deg, *P*_*bootstrap*_ < .001). However, vestibulospinal-lesioned larvae adopted more consistent swim directions through consecutive swim bouts (Figure S2D). They veered less (Figure S2E, 10.68 deg vs. 9.84 deg, *P*_*median-test*_ = .019) and achieved greater depth changes through sequence of bouts compared to sibling controls (Figure S2F, 5.64e-2 deg vs. 0.10 deg, *Pbootstrap* < .001).

We conclude that ascending neurons in the tangential nucleus, and not vestibulospinal neurons, are critical to maintain stable and consistent heading required for effective navigation in depth.

Loss of vestibular function should disrupt posture and locomotor behaviors. We investigated kinematic features that determine swim directions in the vertical axis (Figure S3A). During bouts, larval zebrafish utilize a three-step strategy that allows them to climb and dive while maintaining their preferred horizontal posture^22,43,49^ (Figure S3A, Table 1): First, larvae steer by rotating their their body. Next, they coordinate propulsive forces generated by undulatory thrust and pectoral-fin-based lift. Finally, they rotate back toward their preferred posture. The strength of each of these behaviors can be parameterized as a gain, to indicate the how strongly fish steer (Figure S3B), achieve vertical translocation through lift (Figure S3C), and restore posture (Figure S3D).

Compared to sibling controls, vestibular-impaired larvae exhibited higher steering gain (Figure S3E, *P*_*t-test*_ < .001, *P*_*t-test*_ < .001, *P*_*t-test*_ = .002, Table 2) and lower lifting gain (Figure S3F, *P*_*t-test*_ < .001, *P*_*t-test*_ = .036, *P*_*t-test*_ = .005, for *otogelin* mutants, tangential, and vestibulospinal lesions, respectively). *otogelin* mutants and vestibulospinal lesions resulted in significant decrease in the righting gain (Figure S3G, *P*_*t-test*_ < .001, *P*_*t-test*_ = .180, *P*_*t-test*_ = .001, for *otogelin* mutants, tangential, and vestibulospinal lesions, respectively). Taken together, these data confirm that, as expected, perturbations of vestibular sensation or vestibular neurons disrupt swim kinematics. Specifically, after lesions, larvae can still change depth but they do so with more eccentric posture and poorly coordinated lift (Figure S3H). These results are consistent with the increased variability seen in *otogelin* mutants (Figure 2B), and after lesions of either ascending tangential nucleus neurons (Figure 3C) or vestibulospinal neurons (Figure S2C).

### Gravity-guided stabilization of swim directions for vertical navigation is mediated by the midbrain nucleus INC/nMLF

Ascending neurons of the tangential nucleus project to the INC/nMLF, which sends descending axons to the spinal cord to control locomotion^24,50–53^ (Figure 4A). We reasoned that the INC/nMLF might be the final supraspinal node in a circuit for gravitational control of heading during vertical navigation. We therefore lesioned large descending neurons in the INC/nMLF (Figures S1C, 4A and 4B). Ablation slightly increased directional variability (Figure 4C, 14.29 deg vs. 14.60 deg, controls vs lesions, *P*_*bootstrap*_ = 0.075), and reduced consistency of heading (Figure 4D, Table 2). Similar to effects seen in *otogelin* mutants and after ascending tangential neuron lesions, larvae with INC/nMLF lesions showed increased veering (Figure 4E, 5.98 deg vs. 6.51 deg, *P*_*median-test*_ = .002), and were less efficient at changing depth (Figure 4F, 0.14 vs. 0.11, *P*_*bootstrap*_ < .001).

**Figure 4:**
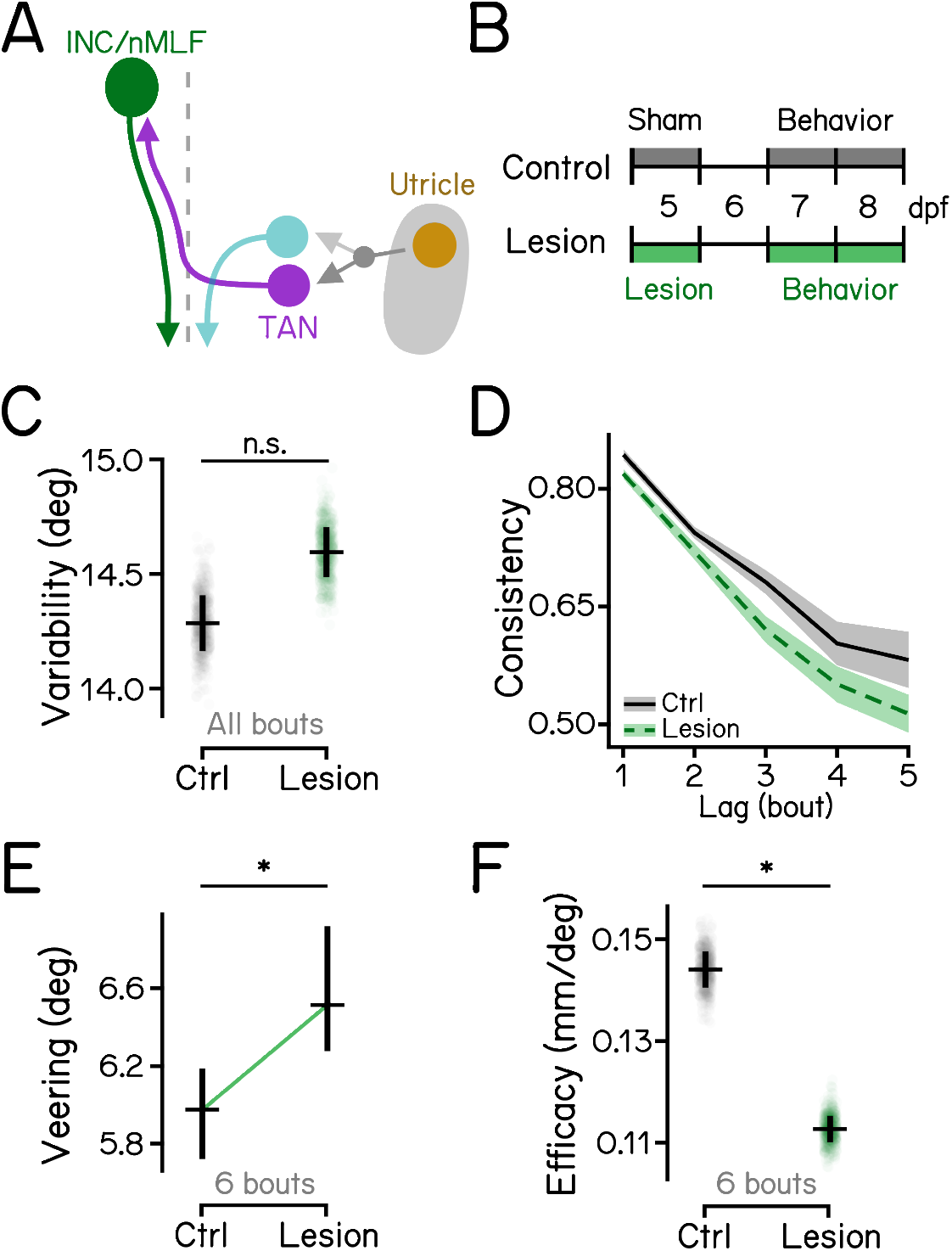
Descending neurons in the INC/nMLF are indispensable for vertical navigation. **(A)** Schematic diagram of the hindbrain-midbrain circuit. Descending neurons in the INC/nMLF (green) receive contralateral vestibular inputs from the tangential nucleus (magenta). **(B)** Experimental diagram of INC/nMLF lesions and behavior assays. Refer to Figure S1C for INC/nMLF lesions. **(C)** Swim direction variability compared between INC/nMLF-lesioned larvae and controls with Boot-strapped standard deviations shown as error bars. n = 18940/22363 bouts from 37/36 fish for controls/lesions. *P*_*bootstrap*_ = 7.53e-2. **(D)** Swim direction consistency plotted as a function of the number of bouts in the sequence. Shaded bands indicate standard deviations of the slope estimated by bootstrapping. **(E)** Veering through 6 consecutive bouts plotted in median with 95% confidence intervals. n = 1215/1340 6-bout series for controls/lesions. *P*_*median-test*_ = 2.13e-3. **(F)** Depth change efficacy plotted with bootstrapped slopes plotted as data points and error bars showing standard deviations. *P*_*bootstrap*_ = 3.00e-13. Abbreviations: TAN, the tangential nucleus; INC, interstitial nucleus of Cajal; nMLF, nucleus of the medial longitudinal fasciculus. See also Table 2 for statistics.

Larvae with lesions of descending neurons in the INC/nMLF recapitulated phenotypes observed in *otogelin* mutants and after lesions of ascending neurons in the tangential vestibular nucleus. Specifically, all three disrupt heading consistency across a series of bouts and show increased veering. Together, these decrease the efficacy of changing depth. Taken together, our results reveal a circuit from the inner ear to the spinal cord responsible for gravitational control of heading during vertical navigation.

## DISCUSSION

We define a circuit that uses gravitational information to control heading for effective vertical navigation. Larvae use a series of swim bouts with consistent heading to change depth in the dark. Loss of either the utricular otoliths, utricle-recipient ascending neurons in the tangential vestibular nucleus, or spinal-projecting neurons in the INC/nMLF all caused fish to swim with more variable heading and excessive veering, leading to inefficient vertical navigation. Taken together, this work reveals ancient brainstem architecture that uses gravitational cues to move effectively through the world.

### Circuit architecture and computations for vertical navigation

Our work argues that the INC/nMLF contributes to vertical navigation. In larval zebrafish, the INC/nMLF is best known for its role in regulating swim posture and speed^50–53^. Further, the tangential-INC/nMLF circuit controls a vestibular-induced body bend reflex that allows fish to maintain posture in the roll (barbecue) axis^24^. Across mammals, INC/nMLF is perhaps best known as the site of the neural integrator for vertical/torsional eye position^54–56^, a computation that acts as a short-term memory for motor commands^57^. Saccades that move the eyes to a new position are instantiated with short bursts of neuronal activity. When integrated, these bursts provide the signal necessary for extraocular motor neurons to maintain muscle tension, stabilizing gaze at the new position. Similarly, integration transforms vestibular representations of head velocity into eye position for a proper vestibulo-ocular reflex^58^.

We propose that the utricle-tangential-INC/nMLF circuit stores and uses a short-term memory of gravity-derived signals for vertical navigation. We observed that larval zebrafish maintain their heading across a series of bouts. The timescale of this phenomena suggests the existence of a short-term memory for commands to shape posture and kinematics. *otogelin* mutants show profoundly disrupted vertical navigation in darkness. Therefore, while heading may persist partly due to inertia, a gravity-derived neural command must contribute as well. Intriguingly, the idea that oculomotor integrator circuits might serve a role in navigation has a recent parallel: in larval zebrafish, the nucleus prepositus hypoglossi (NPH), best known as the neural integrator for horizontal eye movements, may integrate self-motion signals in the yaw plane^41^.

Unlike perturbations to the utricle-tangential-INC/nMLF circuit, larvae without vestibulospinal neurons veer less and navigate more efficiently. While puzzling, this finding is important for two reasons. Firstit demonstrates that not all lesions to the utricle-recipient neurons lead to disrupted navigation. Second, it shows that loss of gravity sensation and lesions of vestibular nuclei perturb swim kinematics similarly, leading to increased variability in bout direction. Since we see both increases and decreases to navigation performance after lesions, we infer that changes to gravity-guided heading can be dissociated from changes to swim kinematics. However, we do not yet know why fish without vestibulospinal neurons veer less. To solve this problem, it will be crucial to define and compare the spinal targets of vestibulospinal neurons^59^ to the targets of descending neurons in the INC/nMLF^52,60^. These findings set the stage to explore the integration of gravity-derived information from direct and indirect brainstem projections to the spinal cord.

### Are larval zebrafish truly navigating depth?

Environmental cues such as light, food, or behavioral state can guide vertical aquatic navigation. Many species of aquatic animals migrate up/down^61^, following the 24-hour cycle of the zooplankton diel vertical migration^62^. Ocean sunfish will perform deep dives during the day to feed in the mesopelagic zone, returning to the surface to warm up^63–65^. Elephant seals dive during sleep to avoid predators^66^. Most pertinently, larval zebrafish can dive/surface following changes to illumination^67–69^ and anxiogenic/anxiolytic drugs^70^, and tend to occupy the top third of the water column in a tall (36 cm) tank^71^.

All behavior in the current study was measured in complete darkness without a defined goal, raising questions of terminology. One proposal would classify the behavior we observe as “gravity-guided orientation,” because larvae seek to arrive at a more preferable depth, rather than navigating to a specific location like a nest^72,73^. However, “orientation” may refer to both a stationary/perceptual response and a locomotor activity in its definition^74^. To avoid ambiguity, we refer to the behavior we see as “gravity-guided navigation,” consistent with a broader view of what comprises navigation^42^. By studying unconstrained vertical navigation, our work sets the foundation for exploration of more complex behavior paradigms. Future work will introduce perturbations, enable goal-directed tasks, and deliver additional stimuli to understand gravity’s influence on navigation.

### Limitations

A potential caveat of our loss-of-function approach is that the lesions were done at different ages. We did not observe regrowth of neuronal cell bodies at any of the lesion sites by the time of behavior assays among the fish we examined, so we do not expect differential regeneration to affect our results. Larvae might exhibit different levels of adaptation to the impairments before behavioral assessment. Given the consistency in effects across constitutive loss (mutants), 4 dpf (tangential) and 5 dpf (INC/nMLF) lesions, we think this explanation is unlikely. Future work with longitudinal behavioral assays will permit investigation of the mechanisms of adaptation and rehabilitation after circuit disruption.

### Conclusion

We define a sensorimotor circuit that uses evolutionarily-conserved brainstem architecture to transform gravitational signals into stable heading for effective vertical navigation. The work lays a circuit-level foundation to understand persistent signals that guide locomotion, and how vestibular inputs allow animals to move efficiently through their environment.

## MATERIALS AND METHODS

### Fish husbandry

All procedures involving larval zebrafish (*Danio rerio*) were approved by the New York University Langone Health Institutional Animal Care & Use Committee (IACUC). Zebrafish embryos and larvae were raised at 28.5°C on a standard 14:10 h light:dark cycle. Larvae were raised at a density of 20-50 in 25-40 ml of E3 medium in 10 cm petri dishes before 5 days post-fertilization (dpf). After 5 dpf, larvae were maintained at densities under 30 larvae per 10 cm petri dish and were fed cultured rotifers (Reed Mariculture) daily.

### Fish lines

Experiments were done using wild type fish with a mixed background of AB, TU, WIK, and SAT. Larvae for lesion experiments were on the mitfa-/- background to remove pigment. Photoablations of ascending neurons in the tangential nucleus were performed in *Tg(−6.7Tru.Hcrtr2:GAL4-VP16,Tg(UAS:EGFP)*^35^. Photoablations of vestibulospinal neurons and neurons of the INC/nMLF were performed on the *Is(nefma:hsp70l-LOXP-GAL4FF),Tg(UAS:EGFP)* background, derived from stl601Tg^75^, henceforth called *Tg(nefma::EGFP). otogelin* mutants were rks^vo66/vo66 44^.

### Vestibular manipulations and photoablations

*otogelin* mutants were screened at 2 dpf for bilateral loss of utricular otoliths. Photoablations of ascending neurons of the tangential nucleus were performed in *Tg(−6.7Tru.Hcrtr2:GAL4-VP16; UAS:EGFP)* larvae at 4 dpf. Lesions of the vestibulospinal neurons and the nMLF were performed in *Tg(nefma::EGFP)* on day 6-7 and 5 dpf, respectively.

All lesions were done using a 2-photon laser as previously described^34^. Briefly, larvae were anesthetized in 0.2 mg/ml MESAB and then mounted in 2% low-melting point agarose. Neurons of interest were identified and imaged using an upright microscope (ThorLabs Bergamo) with an 80 MHz Ti:Sapphire oscillator-based laser at 920 nm (SpectraPhysics MaiTai HP). A separate high-power pulsed infrared laser (SpectraPhysics Spirit 8W) was used for photoablation (1040 nm, 200 kHz repetition rate, 400 fs pulse duration, 1-4 pulses per neuron over 10 ms at 25-75 nJ per pulse). Lesion controls were sibling fish and were anesthetized for comparable durations to lesioned larvae. Lesioned and control sibling larvae were allowed to recover at 28.5°C until behavioral measurements.

### Behavioral measurements

Methods to measure behavior, including apparatus design, hardware, software and procedures, have been extensively detailed^43^. Briefly, larvae at 7 dpf were transferred from petri dishes to behavior chambers. An experimental repeat consisted of a single clutch of larvae run in at least 3 behavioral apparatus. Each apparatus contains 5-8 larvae per standard chamber or 2-3 fish per narrow chamber filled with with 25-30/10-15 ml of E3, respectively. Behavior recordings were started in the morning or around noon on day 7, which is equivalent to circadian time 1-3 or zeitgeber time 1-3, and lasted for approximately 48 hours. After 24 hours of recording, programs were paused for 30 minutes for feeding where 1-2 ml of rotifer culture was added to each chamber. Larvae were removed from the apparatus approximately 48 hours after the start of the experiment. Data during circadian/zeitgeber day were used for all analyses.

### Behavioral analysis

Data was analyzed using our previously published pipeline^43^. In brief, the location of a fish and its pitch axis posture were extracted and saved in real time when a single fish was present in the field of view with its body plane perpendicular to the light path. Data from each experimental repeat were concatenated and swim bouts (defined as a duration where fish swam faster than 5 mm/s) were detected. Swim bouts were aligned at the time of the peak speed for subsequent analysis. Durations between swim bouts where speed was lower than 5 mm/s were considered inter-bout intervals. Bout and fish numbers for each condition are reported in figure legends. Table 1 defines each analysis parameter.

Only consecutive swim bouts were used for autocorrelation analysis, which examined the relationship between bouts with different lags. The lag between two bouts in the same bout series was determined by the number of inter-bout intervals that elapsed in-between. A lag of 1 defines adjacent swim bouts. Bout pairs with different lags were extracted from sequential bouts in a series. For example, a series of 4 consecutive bouts yields 3 pairs of adjacent bouts, 2 pairs of lag-2 bouts, and 1 pair of lag-3 bouts.

### Statistics

All measurements and statistics have been reported in Tables 1 and 2, including expected value, variance, and confidence intervals of parameters. Sample sizes (e.g. number of fish and bout numbers) are included in figure legends. Below we describe the statistical analyses used to compare parameters between conditions.

Median absolute deviation was used to quantify swim direction variability. For variability and consistency, we calculated bootstrapped means for each condition (control and experiment) and took their differences for statistical analysis. Mean and the standard deviation of differences were used to determine the two-tailed P value. For veering, one measurement was calculated from each set of 6-bout sequences. Because veering values are non-parametric, the median test was used to determine significance between two conditions. For depth change efficacy, one fitted slope was calculated from each bootstrapped sample. Difference between two conditions (control and experiment) were calculated and were used for two-tailed significance test. For steering gain, lifting gain, and righting gain, one value was calculated from each experimental repeat for unpaired two-tailed t-test.

### Data Availability

All raw data are available at the Open Science Framework DOI: 10.17605/OSF.IO/AER9F

### Code Availability

All analysis code is available at the Open Science Framework DOI: 10.17605/OSF.IO/AER9F

## ACKNOWLEDGMENTS

Research was supported by the National Institute on Deafness and Communication Disorders of the National Institutes of Health under award numbers R01DC017489 (DS), and F31DC019554 (KRH), and the National Institute of Neurological Disorders and Stroke under award numbers, T32NS086750 (KRH) and R61NS125280 (DS), by the Leon Levy Foundation (YZ), and the Rainwater Charitable Foundation (YZ). The authors thank Christina May and other members of the Schoppik and Nagel laboratories for their valuable feedback and discussions.

## AUTHOR CONTRIBUTIONS

Conceptualization: YZ and DS. Methodology: YZ. Investigation: YZ, HG, FA, KRH, DEE. Formal Analysis: YZ. Visualization: YZ. Writing: YZ and DS. Editing: YZ and DS. Supervision: DS. Funding Acquisition: YZ, KRH, and DS.

## AUTHOR COMPETING INTERESTS

The authors declare no competing interests.

---

**Figure S1:**
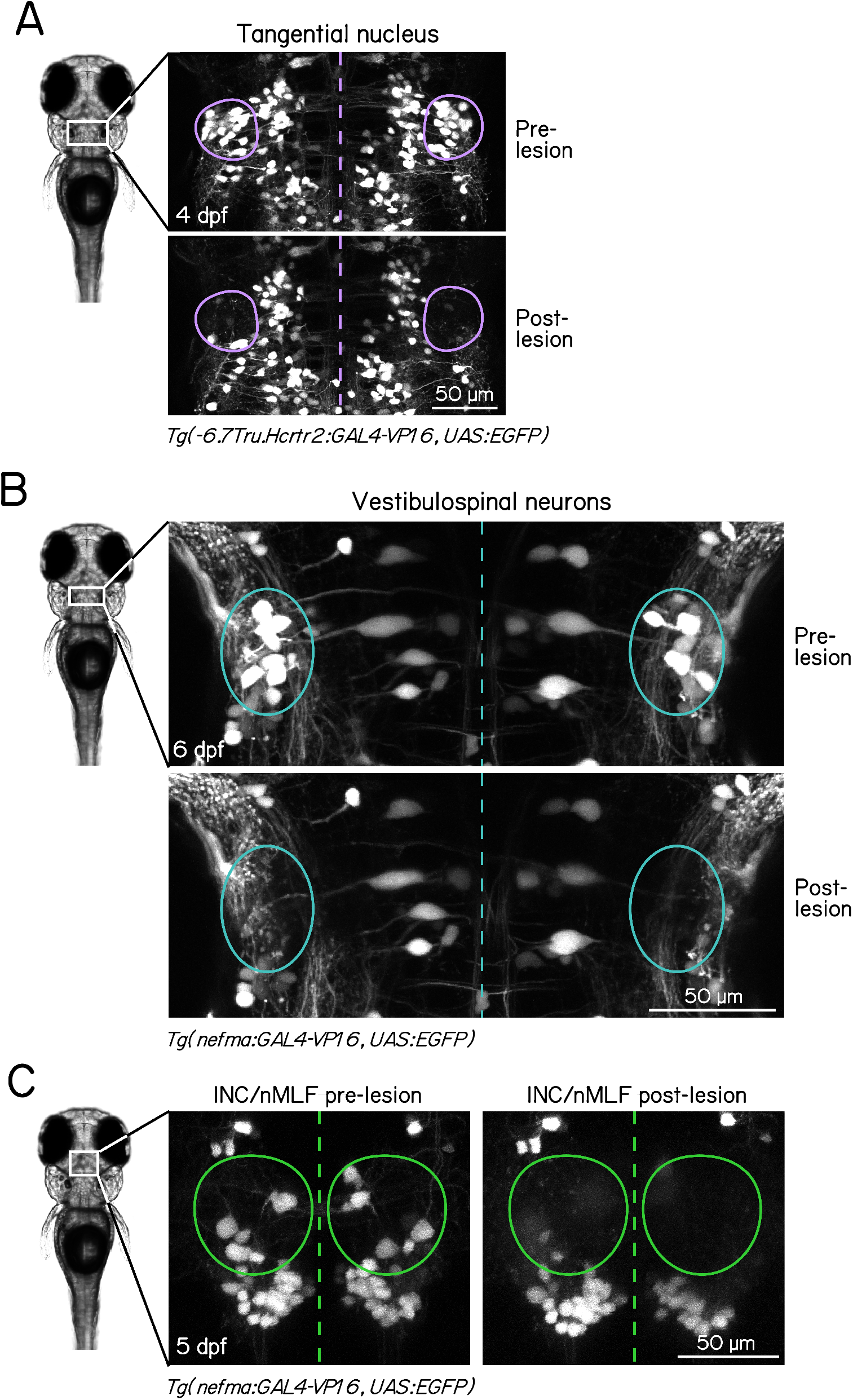
Example larvae before and after photoablation. **(A)** Before and after lesions of the tangential vestibular nucleus (circled) in a 4 dpf larvae. Scale bar: 50 µm. **(B)** Before and after lesions of the vestibulospinal nucleus (circled) in a 6 dpf larvae. Scale bar: 50 µm. **(C)** Before and after lesions of large neurons in the interstitial nucleus of Cajal/the nucleus of the medial longitudinal fasciculus (INC/nMLF, circled) in a 5 dpf larvae. Scale bar: 50 µm.

**Figure S2:**
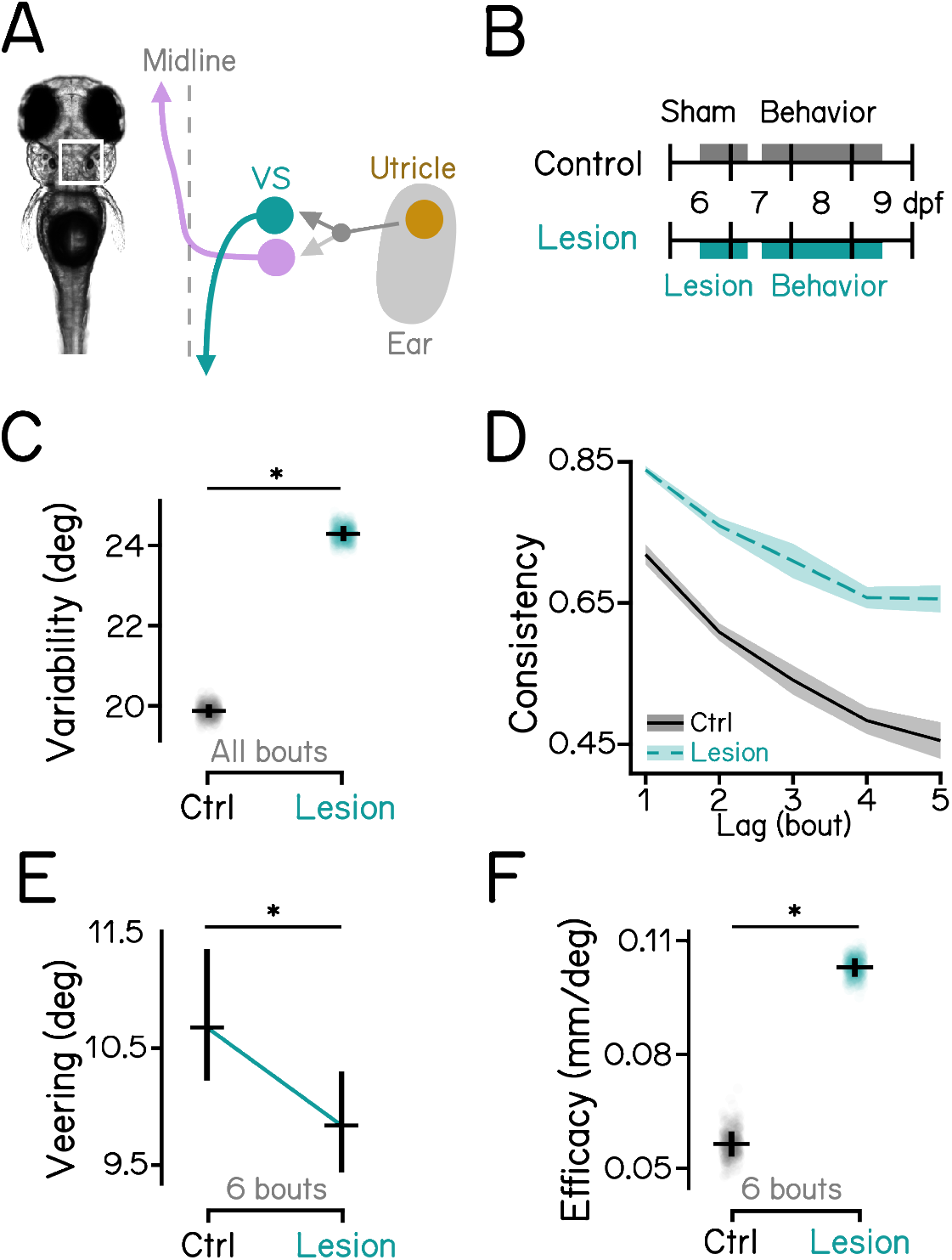
Lesions of vestibulospinal neurons increase postural variability but stabilize veering, improving vertical navigation. **(A)** Schematic view of the inner-ear utricular otolith and the vestibular pathways in the hindbrain of zebrafish. Utricle: utricular otoliths (yellow); VS: vestibulospinal neurons (cyan). **(B)** Diagrams of experimental procedures for lesions of the vestibulospinal nucleus and behavioral assays. See Figure S1B for examples of lesions. **(C)** Swim direction variability compared between vestibulospinal-lesioned larvae and controls. Bootstrapped MAD are plotted as data points with error bars showing standard deviations. n = 18106/17758 bouts from 79/97 fish for controls/lesions. *P*_*bootstrap*_ = 3.87e-70. **(D)** Swim direction consistency plotted as a function of the number of bouts in the sequence. Shaded bands indicate standard deviations of the slope estimated using bootstrapping. **(E)** Veering through 6 consecutive bouts plotted in median with 95% confidence intervals. n = 1471/1076 6-bout series for controls/lesions. *P*_*median-test*_ = 1.88e-2. **(F)** Depth change efficacy plotted with bootstrapped slopes plotted as data points and error bars showing standard deviations. *P*_*bootstrap*_ = 1.17e-27. See also Table 2 for statistics.

**Figure S3:**
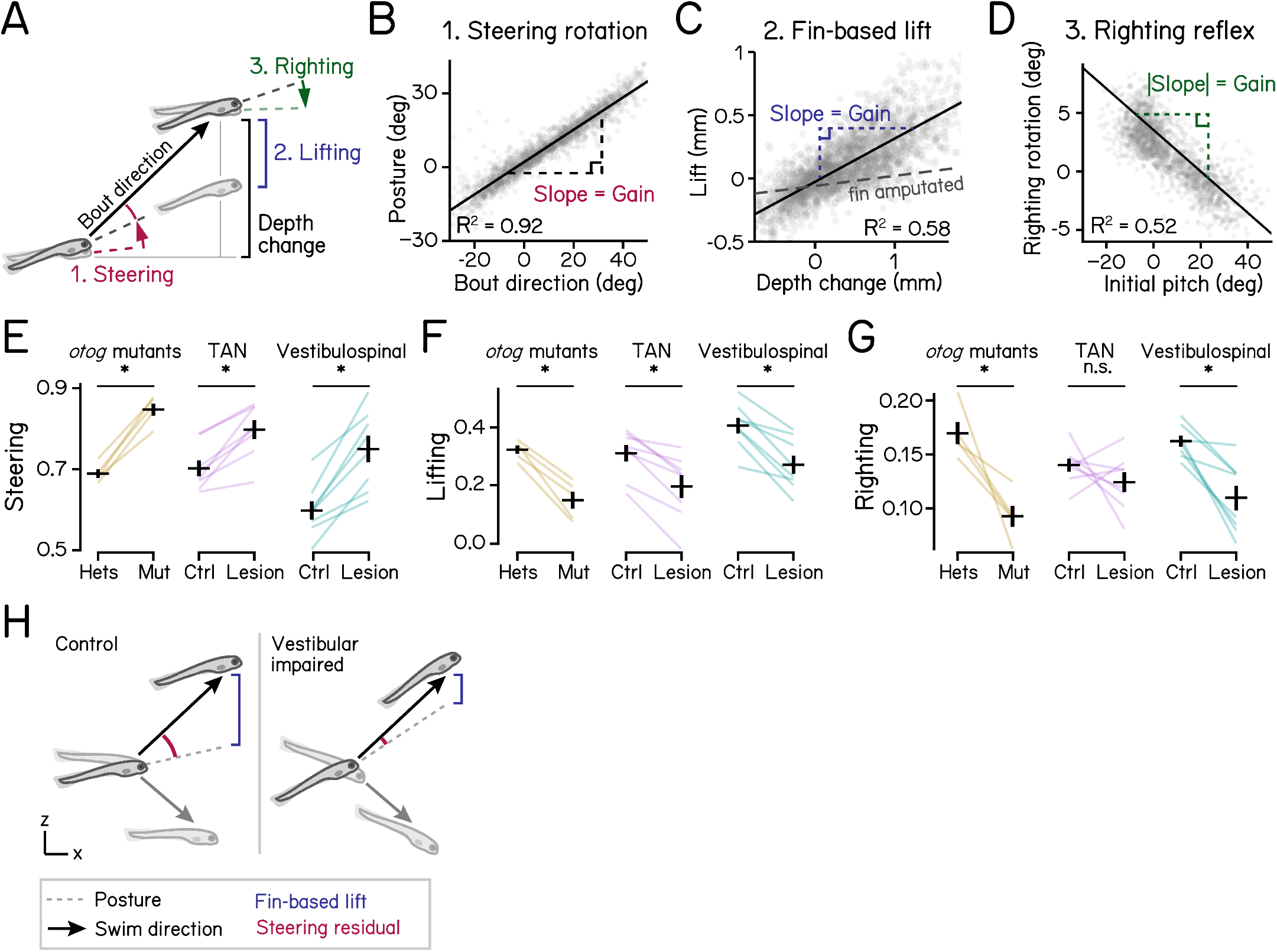
Vestibular contribution to swim kinematics. **(A)** Schematic diagram showing steering, lifting, and righting during a swim bout. Larvae steer toward targeted direction during acceleration (red arrow), use pectoral fins to assist in depth changes (blue), and restore posture to horizontal during deceleration (green arrow). Z displacement generated by lifting (blue) is estimated by subtracting theoretical displacement in depth, calculated from the head direction and x distance, from the total depth change. **(B)** Steering gain is defined as the slope of the best fit line of posture at the time of the peak speed vs. swim direction. n = 121,979 bouts from 537 fish. **(C)** Lift gain is defined as the slope of the best fit line of estimated lift vs. depth change of the swim bout. Pectoral-fin amputation reduces lift (dashed line). n = 33,491/28,604 bouts from 74/78 fish for control/fin-amputated. **(D)** Righting gain is defined as the numeric inversion of the slope of the best fit line of rotation during deceleration vs. initial posture. n = 121,979 bouts from 537 fish. **(E)** Steering gain of vestibular-impaired larvae vs. controls. *otog* mutation: *P*_*t-test*_ =2.33e-5; tangential lesions: *P*_*t-test*_ = 7.41e-3; vestibulospinal lesions: *P*_*t-test*_ = 2.32e-3. N = 5/8/8 experimental repeats for *otog*/tangential lesions/ vestibulospinal lesions. Same as follows. **(F)** Lifting gain of vestibular-impaired larvae vs. controls. *otog* mutation: *P*_*t-test*_ = 7.09e-4; tangential lesions: *P*_*t-test*_ = 3.57e-2; vestibulospinal lesions: *P*_*t-test*_ = 5.90e-3. **(G)** Righting gain of vestibular-impaired larvae vs. controls. *otog* mutation: *P*_*t-test*_ = 6.76e-4; tangential lesions: *P*_*t-test*_ = .180; vestibulospinal lesions: *P*_*t-test*_ = 1.00e-3. **(H)** Summary of effects of vestibular perturbations on bout kinematics. Vestibular-impaired fish swim with more eccentric posture and less fin-based lift. Abbreviation: TAN, tangential. See also Table 1 for parameter definitions and Table 2 for statistics.

## Notes

https://osf.io/aer9f/

### Competing Interest Statement

The authors have declared no competing interest.

